# Reactive microglia fail to respond to environmental damage signals in a viral-induced mouse model of temporal lobe epilepsy

**DOI:** 10.1101/2024.03.06.583768

**Authors:** Glenna J. Wallis, Laura A. Bell, John N. Wagner, Lauren Buxton, Lakshmini Balachandar, Karen S. Wilcox

## Abstract

Microglia are highly adaptable innate immune cells that rapidly respond to damage signals in the brain through adoption of a reactive phenotype and production of defensive inflammatory cytokines. Microglia express a distinct transcriptome, encoding receptors that allow them to dynamically respond to pathogens, damage signals, and cellular debris. Expression of one such receptor, the microglia-specific purinergic receptor *P2ry12*, is known to be downregulated in reactive microglia. Here, we explore the microglial response to purinergic damage signals in reactive microglia in the TMEV mouse model of viral brain infection and temporal lobe epilepsy. Using two-photon calcium imaging in acute hippocampal brain slices, we found that the ability of microglia to detect damage signals, engage calcium signaling pathways, and chemoattract towards laser-induced tissue damage was dramatically reduced during the peak period of seizures, cytokine production, and infection. Using combined RNAscope *in situ* hybridization and immunohistochemistry, we found that during this same stage of heightened infection and seizures, microglial *P2ry12* expression was reduced, while the pro-inflammatory cytokine *TNF-a* expression was upregulated in microglia, suggesting that the depressed ability of microglia to respond to new damage signals via *P2ry12* occurs during the time when local elevated cytokine production contributes to seizure generation following infection. Therefore, changes in microglial purinergic receptors during infection likely limit the ability of reactive microglia to respond to new threats in the CNS and locally contain the scale of the innate immune response in the brain.

## Introduction

Viral encephalitis is often accompanied by acute seizures and an increased probability of developing long-term epilepsy. Mice infected intracerebrally with the Daniel strain of Theiler’s Murine Encephalomyelitis Virus (TMEV) develop acute seizures from 3-8 days post-infection (DPI) and exhibit pathologic changes such as reactive glia (microglia, astrocytes, and NG2-glia), infiltration of peripheral immune cells, increased oxidative stress, elevated cytokine production, substantial neuronal degeneration and scar formation in CA1, and increased excitatory synaptic transmission in CA3 of the hippocampus (Libbey *et al*., 2008; Stewart *et al*., 2010; Smeal *et al*., 2012; Loewen *et al*., 2016; Patel *et al*., 2017; Bell, Wallis and Wilcox, 2020; DePaula-Silva *et al*., 2021; Lawley *et al*., 2022). Importantly, mice survive the infection, clear the virus from the brain by 14 DPI, and develop chronic spontaneous seizures as well as cognitive impairments and anxiety-like symptoms commonly associated with human temporal lobe epilepsy (TLE) (Stewart *et al*., 2010; Libbey *et al*., 2011b; Umpierre *et al*., 2014; Patel *et al*., 2017). This animal model of infection-induced TLE provides an opportunity to understand disease mechanisms in the quest to find innovative therapies for the prevention of epilepsy.

The robust scale and extended duration of the pro-inflammatory response may contribute to seizure development following TMEV infection. As the resident immune cells of the brain, microglia rapidly detect pathogens and damage, extend cellular processes, release free radicals locally, and phagocytose dead or damaged cells. Microglia also release cytokines to promote local cell defense mechanisms and release chemokines to recruit peripheral immune cells into the central nervous system (CNS). Finally, microglia can adopt a reactive phenotype to further enhance pro-inflammatory roles (Damisah *et al*., 2020; Henning *et al*., 2023). Two pro-inflammatory cytokines, tumor necrosis factor alpha (TNF-α) and interleukin 6 (IL-6), produced by microglia and macrophages, significantly contribute to acute seizure severity following infection with TMEV (Libbey *et al*., 2011a; Patel *et al*., 2017; DePaula-Silva *et al*., 2021). These cytokines can act directly on neurons to induce necrosis or to promote synaptic scaling in addition to amplifying the immune response (Beattie, Ferguson and Bresnahan, 2010; Stellwagen, 2011; Patel *et al*., 2017; Kano *et al*., 2019; Henning *et al*., 2023). Later, CNS-infiltrating T-cells signal to resolve microglia activation and restore normal cellular function (Gordon and Taylor, 2005; Town, Nikolic and Tan, 2005). However, the escalating damage from the virus, cell death, reactive oxygen species (ROS), increased cytokine expression, and ongoing acute seizures are thought to sustain microglia reactivity and may prolong and amplify pro-inflammatory actions following infection.

Microglia use rapid-acting intracellular calcium signals and kinase/phosphatase pathways to respond to neuronal activity and damage signals (Eichhoff, Brawek and Garaschuk, 2011; Tvrdik and Kalani, 2017; Liu *et al*., 2019; Damisah *et al*., 2020; Hughes and Appel, 2020; Hu, Shi and Gao, 2020; Umpierre *et al*., 2020; Umpierre and Wu, 2021; Umpierre *et al*., 2023). Surveillance by microglia in the healthy brain includes low-level continuous process movements and a low frequency of spontaneous calcium events, whereas damage responses include a robust calcium response and directional process movement (Eichhoff and Garaschuk, 2011; Brawek and Garaschuk, 2014; Pozner *et al*., 2015; Bennett *et al*., 2016; Brawek *et al*., 2017a; Hughes and Appel, 2020; Umpierre *et al*., 2020). Reactive microglia undergo morphological changes which can include a more ameboid appearance with thickened major processes, retraction of fine processes (Stence, Waite and Dailey, 2001), and an altered expression profile of surveillance genes associated with cytoskeleton reorganization and damage signal recognition (Bennett *et al*., 2016; Srinivasan *et al*., 2016; Lively and Schlichter, 2018; DePaula-Silva *et al*., 2019; Hammond *et al*., 2019). Yet, in most acute models of inflammation, microglia generally display elevated surveillance rates and calcium-mediated damage responses. After inflammatory events, microglia have been reported to have increased spontaneous calcium activity (Pozner *et al*., 2015). Some have reported increased process movement at acute time points (Orr *et al*., 2009; Eyo *et al*., 2014; Avignone *et al*., 2015; Pozner *et al*., 2015; Riester *et al*., 2020), and either reduced or elevated process movements at later time points (Brawek and Garaschuk, 2014; Gyoneva *et al*., 2014). In addition, reactive microglia in the TMEV model have been reported to have significant gene expression changes in damage sensing receptors such as P2YR12 (DePaula-Silva *et al*., 2019). However, the ability of microglia to sense and respond to damage signals during this period has not been investigated. To determine if microglia damage responses and engagement of calcium signaling are disrupted following TMEV infection, we used acute brain slices obtained from TMEV-infected mice expressing tdTomato (TdT) and the fluorescent calcium sensor GCaMP5G in microglia. We applied an automated signal detection algorithm that consistently identified regions of calcium activity within microglia, and we report the spontaneous activity characteristics, as well as those induced by laser burn and exogenous adenosine triphosphate (ATP) application, for both microglia somas and processes. We have identified functional deficits in microglial detection of cellular damage and purinergic signals during the acute TMEV-infection period, and this may disrupt actin-dependent movements normally observed in the processes of microglia in the resting state. We also identified heterogeneity in the ability of microglia to detect specific damage signals and in the ability to transmit calcium signals through different subcellular regions (soma *versus* processes). Thus, hippocampal microglia, during the peak of the pro-inflammatory response after TMEV infection, may be less responsive to escalating damage. The present work demonstrates that a better understanding of the fundamental interactions between microglia and their environment will allow us to identify new ways to intervene in neuroinflammatory conditions.

## Results

### TMEV infection induces seizures during the acute infection period

Mice heterozygous for Cx3cr1-EYFP-creERT2 and PC::G5-tdT were administered tamoxifen (TAM) to induce recombination and allow expression of the genetically encoded calcium sensor GCaMP5G (G5) and cytosolic tdTomato (TdT) in microglia. By including both fluorophores in microglia, we were able to track the position of the somas and processes of cells (TdT) and identify cellular regions where calcium signaling was engaged (G5). By delaying experiments until after 35 days from TAM injection (**Figure 1A**), newly differentiated peripheral macrophages infiltrating into the CNS display only EYFP, making them distinguishable from microglia which express both TdT and G5 (Parkhurst *et al*., 2013). In TMEV infected mice, and as previously described (DePaula-Silva *et al*., 2021; Batot *et al*., 2022), handling-induced seizures begin around 3 DPI and progress in severity, as measured by a modified Racine scale, through 7 DPI (**Figure 1B**). While mice at 2 DPI do not present with seizures, they do display mild weight loss which continues throughout the acute seizure period (**Figure 1C**). The number of seizures per day peaked at 5 DPI, and it has been previously reported that pro-inflammatory cytokine and ROS levels are also highly elevated at 5 DPI (Bhuyan *et al*., 2015; Patel *et al*., 2017). None of the control / phosphate-buffered saline (PBS)-injected mice exhibited seizures.

**Fig. 1.**
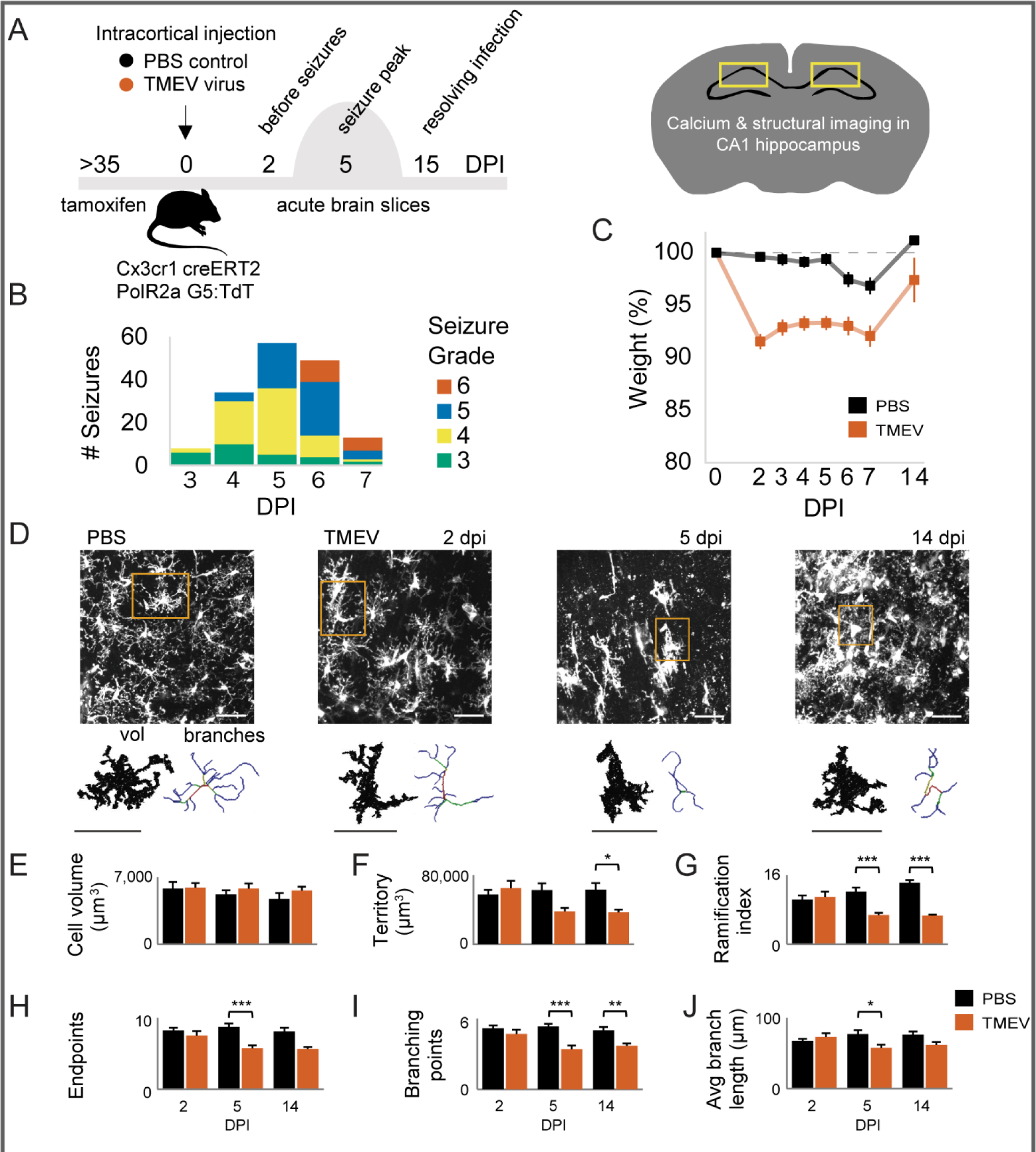
3D morphology of microglia after viral encephalitis display fewer branch ramifications but similar intracellular volume. **(A)** Timeline of experiment. Mice heterozygous for Cx3cr1-EYFP-cerERT2:PC-GCaMP5G were administered i.p. tamoxifen to induce expression of the calcium indicator GCaMP5G (G5) and red TdTomato (TdT). After more than 35 days, mice were injected intracranially with TMEV (test group) or PBS (control), were monitored for seizures twice a day from 3-7 days post-infection, and acute brain slices were prepared at 2, 5, and 15 days post-injection (DPI). **(B)** Mild handling-induced seizures manifest at 3 DPI and progress to more severe grade seizures through 7 DPI. PBS-injected mice did not have seizures. **(C)** TMEV-infected mice have weight loss. **(D)** Morphology of microglia in maximum 81 µm image projections. Cell volume is 3D model of fluorescent image, and cell territory is 3D shape that includes the branch endpoints. The 3D morphologic features of microglia were measured with a semi-automated 3DMorph script that determines the best-fit branching skeleton inside the fluorescent cell volume. Measurements were acquired for **(E)** cell volume, **(F)** cell territory, **(G)** ramification index (the ratio of territory to volume), **(H)** endpoints (measured as the most distant pixel on a branch path), **(I)** number of branch points for each cell territory, and **(J)** the average branch length. PBS-injected mice n=7, 6, & 8 and TMEV-injected mice n=6, 6, & 7 at 2, 5, & 14 DPI respectively. 2-way ANOVA with Bonferroni’s test p<*0.05, **0.01, ***0.001. Scale bar = 50 µm.

### Altered microglia 3D morphology following TMEV infection

Microglia respond to brain infection by transitioning to a reactive phenotype with heightened immune functions and more ameboid morphology with short thick processes (Bennett *et al*., 2016; Srinivasan *et al*., 2016; Lively and Schlichter, 2018; DePaula-Silva *et al*., 2019; Lawley *et al*., 2022). To determine the timing and extent of microglia structural changes after TMEV brain infection in live tissue, we imaged TdT-expressing microglia within acutely prepared hippocampal brain slices. We focused on this region as TMEV infects primarily pyramidal neurons, causing significant death in CA1/CA2, and forming a likely seizure onset zone in the hippocampus (Libbey *et al*., 2008; Patel *et al*., 2017). Microglia structure was measured using the semi-automatic 3DMorph (York *et al*., 2018) on one Z-stack per mouse, with an average of 23 microglia identified per stack in PBS images and an average of 14 microglia identified per stack in TMEV images at 2 DPI, 11 microglia per stack in TMEV images at 5 DPI, and 35 microglia per stack in TMEV images at 14 DPI (**Figure 1D-J**).

At each time point following TMEV-infection, six parameters of microglia cell size and branch ramification were compared to microglia from control (PBS-injected) mice (**Figure 1E-J**). The microglia parameters from mice treated with PBS were consistent with previously reported microglia measurements both *in vivo* and *ex vivo* (York *et al*., 2018). First, the cell volume was not significantly different at any timepoint for microglia in acute brain slices from TMEV-infected mice (**Figure 1E**). Second, the cellular territory was significantly reduced for microglia in acute brain slices from TMEV-infected mice (**Figure 1F**; mean±SEM: 36,814±3,295 µm^3^, p<0.05). Third, the ramification index (ratio of territory to cell volume) was decreased at both 5 and 14 DPI in slices from TMEV-infected mice (**Figure 1G**; 6.8±0.5 and 6.7±0.2, p<0.001). This finding is likely because PBS microglia have thin but extensively branched processes that increase their overall cell volume, while microglia from TMEV-infected mice have shorter, thicker branches which decreases their territory, so overall volume was not changed. Finally, the number of process endpoints was decreased at 5 DPI (**Figure 1H**; 5.7±0.4, p<0.001), the number of branching forks was decreased at 5 and 14 DPI (**Figure 1I**; 3.4±0.3, p<0.001 and 3.7±0.2, p<0.01), and the average length of a branch was shorter at 5 DPI (**Figure 1J**; 58±5 µm, p<0.05). Thus, our 3D microglia assessment demonstrates reduced branch ramification with a similar microglia intracellular volume for two weeks following viral brain infection and expands upon previous findings of microglial hypertrophy in the TMEV-model (Loewen *et al*., 2016; Bell, Wallis and Wilcox, 2020). Importantly, less ramified microglia likely have fewer contacts with synapses, vasculature, and other brain cells within the same territory following brain infection.

### Frequency of spontaneous calcium activity in microglia is decreased following TMEV infection

Microglial fine process extension and retraction during continuous surveillance activity is coupled to actin-cytoskeleton reorganization via intracellular calcium transients, NADH availability, and potassium currents (Dissing-Olesen *et al*., 2014; Swiatkowski *et al*., 2016; Bernier *et al*., 2019; Franco-Bocanegra *et al*., 2019). To determine if TMEV infection affects the number and frequency of spontaneous calcium events in microglia regions of interest (ROIs)a, we used 2-photon imaging for 15-minute epochs in acute hippocampal brain slices obtained from either PBS or TMEV-injected mice (**Figure 2A**). A low rate of spontaneous calcium events was detected in microglia from PBS-injected mice (**Figure 2A-D**; 0.295±0.100 events/min). Similarly, low rates of spontaneous calcium transients have been reported with different calcium detection methods (Eichhoff, Brawek and Garaschuk, 2011; Brawek and Garaschuk, 2014; Del Moral *et al*., 2019; Umpierre *et al*., 2020).

**Fig. 2.**
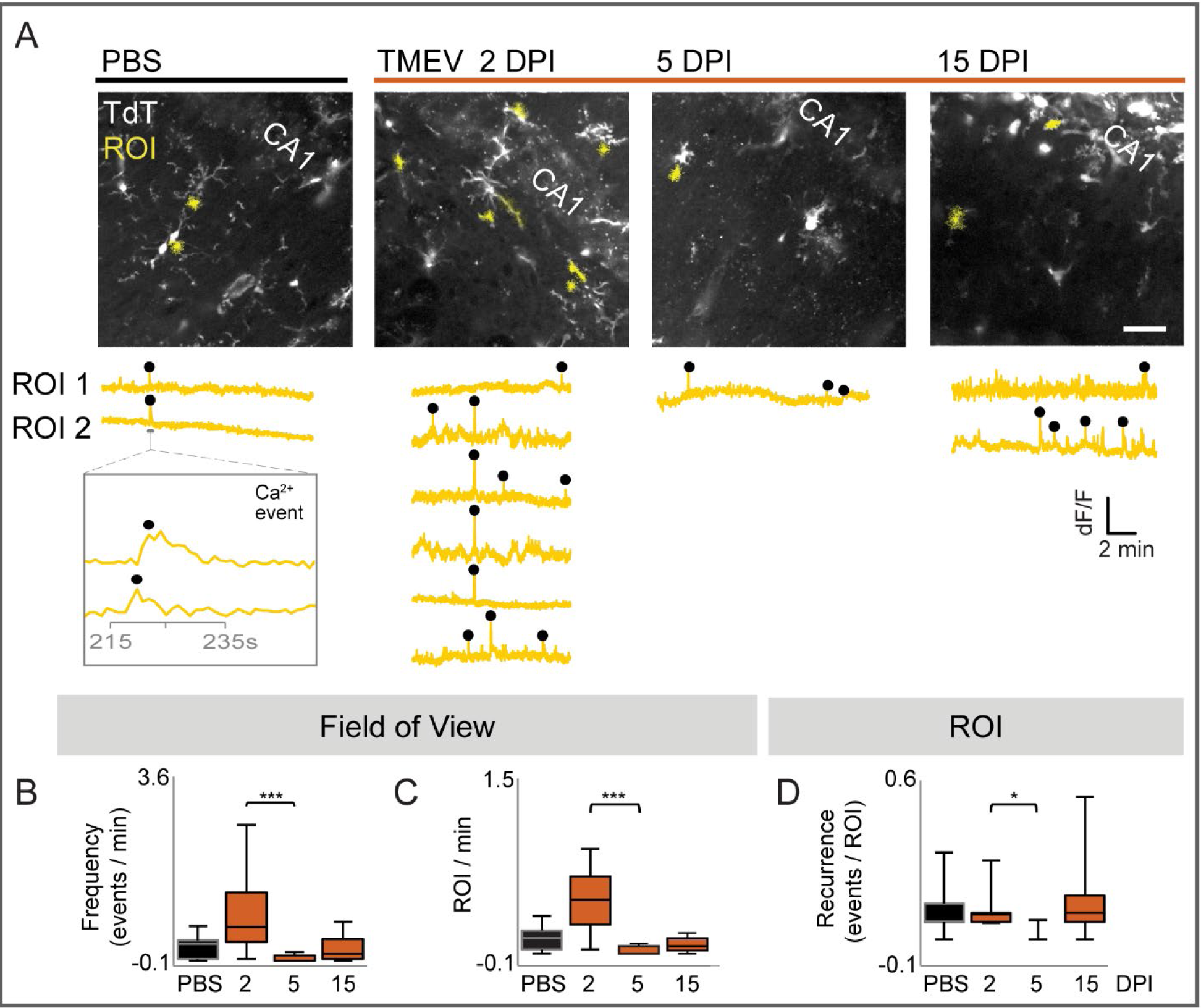
Spontaneous calcium events and active calcium regions in microglia from acute brain slices are reduced at 5 DPI. **(A)** Acute brain slices were recorded for approximately 15 minutes for microglial spontaneous calcium events in CA1 hippocampus. Maximum-over-time images of microglia with regions of interest (yellow) detected as changes in G5 fluorescence using Suite2p. G5 dF/F traces with calcium events (black dot) identified with the Matlab peak finder for each ROI. **(B)** The frequency and **(C)** number of active ROIs, and **(D)** the recurrence (number of events occurring in the same ROI) normalized per min for n=FOV/mice, n=9/8, 13/8, 9/8, & 7/7 from PBS (2-16 DPI), TMEV (2 DPI), TMEV (4-6 DPI), and TMEV (14-16 DPI) respectively. Kruskal-Wallis test with Dunn’s multiple comparison *p<0.05, **p<0.01, ***p<0.001. Scale bar = 50 µm.

In brain slices obtained from TMEV-infected mice, there was no significant change in the frequency of microglial calcium transients at 2 DPI compared to those imaged in PBS-treated mice. However, there was a significant reduction in the frequency of calcium transients at 5 DPI compared to 2 DPI (**Figure 2B**; 5 DPI 0.048±0.029 events/min, p<0.001). The frequency of spontaneous events at 15 DPI was not significantly different from PBS (**Figure 2B**), indicating that frequency of spontaneous calcium activity had returned to baseline levels following viral clearance. The number of calcium events occurring in the same ROI (recurrence) over the time of image acquisition (#calcium events/ROI) was also significantly decreased at 5 DPI compared to 2 DPI (**Figure 2D**, p<0.05). Altogether, in the days immediately following viral brain infection, hippocampal microglia display a decrease in spontaneous calcium transients during the peak of acute seizures in the TMEV model.

### Microglia process movement towards tissue damage is impaired following TMEV infection

A high-power laser can burn small regions of brain tissue (Eichhoff, Brawek and Garaschuk, 2011; Brawek *et al*., 2017a), and microglia have a diverse range of membrane receptors that detect cellular fragments and debris released by necrotic cells (Färber and Kettenmann, 2006). Following laser burn, a characteristic fluorescent lesion allows monitoring of the size of tissue damage and the ability of microglia processes to converge on the lesion (**Figure 3A**).To determine whether microglia are capable of responding to additional neurological insult and tissue damage during and after TMEV infection, we evaluated the area of encroachment of microglial processes to laser burns in the CA1 region of hippocampal brain slices obtained from TMEV-infected mice at 2, 5, and 15 DPI. In slices from uninfected PBS-treated mice, microglia on all sides of the burn send their processes toward the burn region and occupy 19.6±1.6% of the central region by the end of a 30-minute imaging session (**Figure 3A**). This is consistent with previous findings that microglia in healthy brain tissue efficiently surround laser damage (Haynes *et al*., 2006; Gyoneva *et al*., 2014). At 2 days after TMEV infection, microglia processes still migrated to the burned area to a similar degree (**Figure 3B**). However, microglia at 5 and 15 DPI failed to mount a complete containment response (**Figure 3B**, 9.5±1.9% and 8.5±0.9% respectively, p<0.001). Thus, activated microglia in the hippocampus of TMEV-infected mice have an impaired ability to send processes towards regions of newly damaged tissue.

**Fig. 3.**
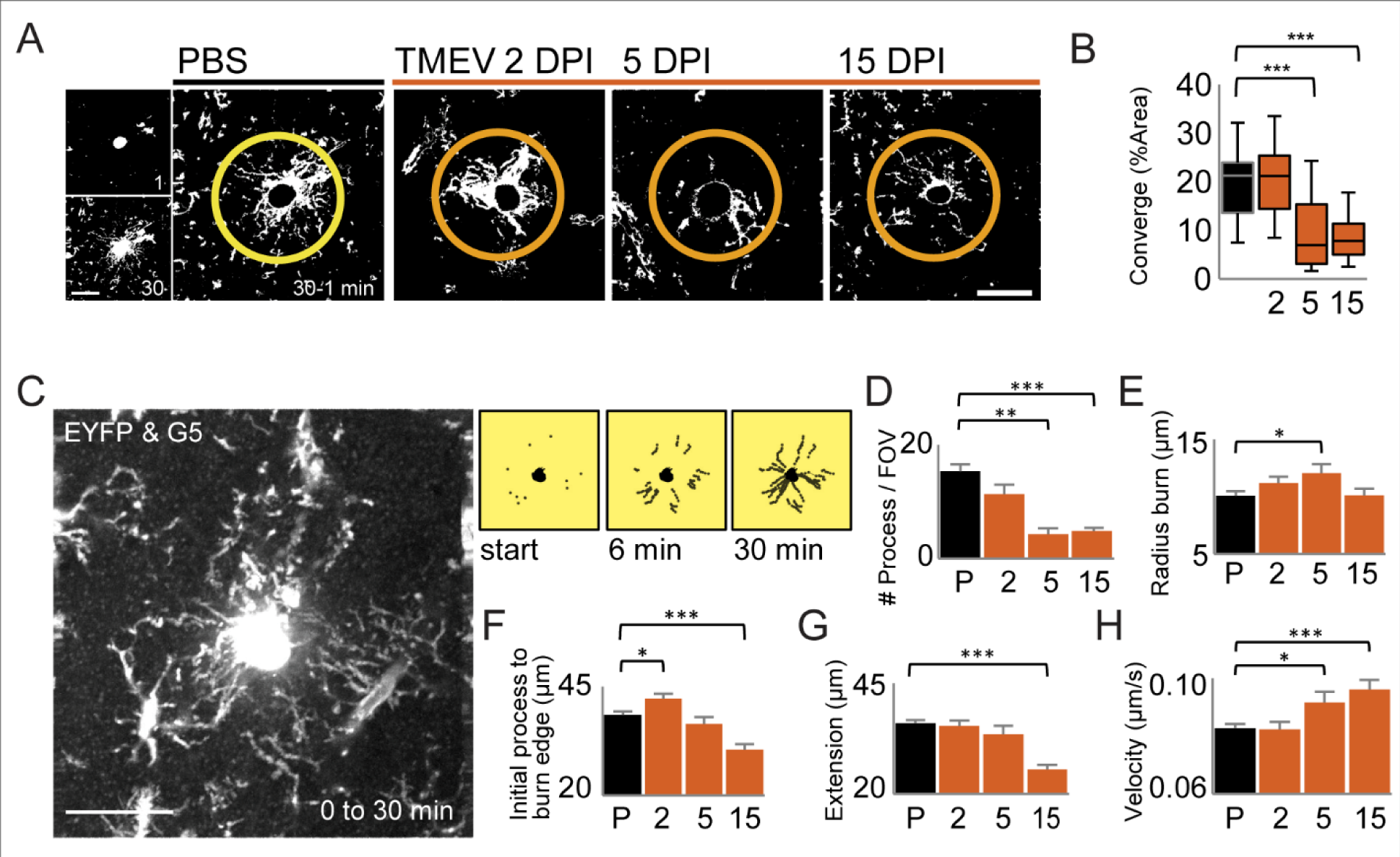
Microglia process convergence around burn damage is reduced following TMEV infection. Microglia processes detect laser burn damage in acute brain slice and respond by sending the processes toward the central burn region. **(A)** The maximum-over-time projections for the starting process location and end location after 30 min were thresholded and subtracted to calculate the area of process convergence around the central burn. **(B)** The median percent area in the central circle was reported with boxes as 75%/25% and error bars 95%/5% for n=mice/images, n=8/27, 7/23, 6/20, 8/30 respectively for PBS and TMEV 2, 5, and 15 DPI. Statistical significance with One-Way ANOVA and Bonferroni’s multiple comparison p<0.001. Scale bar 50 µm. **(C)** Process location was marked every 62 s over 30 minutes and **(D)** the number of processes per field of view (FOV), **(E)** the radius of the burn lesion, **(F)** the initial distance of processes to burn edge, **(G)** the distance of process extension, and **(H)** the velocity of extension were measured for n=mice/images, n=8/19, 5/8, 5/9 & 6/11 from PBS, TMEV 2, 5 and 15 DPI respectively. T-test *p<0.05, **p<0.01, ***p<0.001. Scale bar 50 µm.

Microglia process extension toward the burn was manually tracked every minute for the 30-minute observation period, and the percentage of coverage, number, distance, and velocity of microglia process extension was compared between slices from TMEV and PBS-injected mice (**Figure 3D-H**). At 5 DPI, the same 24 µm^2^ size laser exposure created significantly larger burn lesions in TMEV-infected mice, suggesting that brain tissue during peak seizures and infection is particularly more susceptible to new damage (**Figure 3E**). Upon initiating movement towards the burn, microglia processes were initially 38.6±0.8 µm away from the burn zone in slices obtained from PBS-injected mice. This initial response radius was further away for microglia at 2 DPI (**Figure 3F**; 42.2±1.2 µm, p=0.016). While the average response distance for 5 DPI processes was equivalent to PBS, processes were initially closer to the burn lesion at 15 DPI (**Figure 3F**; 30.5±1.3 µm, p<0.001). To determine the cumulative movement of all processes surrounding the burn, we measured the total process extension within the 30-minute imaging period and found significantly reduced distance of process extension for microglia from 15 DPI TMEV mice (**Figure 3G**; 25.6±1.0 µm, p<0.001). Between 94-100% of tracked processes in all groups had sufficient time to extend and contact the burn zone during the 30-minute imaging period. Most processes appeared to take the shortest path toward the burn, while other processes were clearly seen for several minutes, then appeared to leave the plane of focus, likely to navigate around other cells within the neuropil. Despite fewer microglia processes extending at 5 and 15 DPI, those that did extend had a significantly faster growth velocity **(Figure 3H**; 0.092±0.004 µm/s, p=0.02; 0.096±0.003 µm/s, p<0.001). Together, these data demonstrate that while activated microglia extend fewer processes towards damage following TMEV-infection and seizures, processes that are extended move at a heightened velocity (**Figure 3D,H**). Previously, primed or reactive microglia from a APP23PS45 Alzheimer’s model also displayed faster encroachment on an ATP pipet than those of wild type mice (Brawek and Garaschuk, 2014). Additionally, these data suggest that the reduced coverage by microglia processes around the laser burn in slices from TMEV treated mice is due to fewer numbers of processes being extended toward the burn (**Figure 3A,B**).

### Microglia display three distinct phases of intracellular calcium events following laser burn-induced tissue damage

Three readily observable microglia calcium signaling events have been observed after tissue damage and have previously been reported. First, microglia display discrete intracellular calcium events after neuron puncture, laser burns, or ATP application (Eichhoff, Brawek and Garaschuk, 2011; Brawek *et al*., 2017b). Second, microglia cellular processes begin extending toward damaged neurons or a source of ATP/ADP in a calcium-mediated and actin-dependent manner. Third, microglia processes contact or encase the damaged region (Hines *et al*., 2009; Damisah *et al*., 2020). In the present set of experiments, we induced a laser burn in the hippocampus and tracked microglia process extension with EYFP and G5 over the 30-minute observation period, first under control conditions in slices from PBS-injected mice (**Figure 4**), then compared to slices prepared over the time course of TMEV infection (**Figure 5**). Using the Suite2p toolbox for cell detection and signal extraction, we identified calcium ROIs that contained fluorescent changes (Pachitariu *et al*., 2016). We used local prominence criterion (MATLAB) to identify discrete calcium events (black circles) over the baseline (**Figure 4A**). To correlate subcellular regions with calcium activity, an experimenter overlayed calcium ROIs onto each process extension track, classified the subcellular location, and removed spatially redundant track/ROI pairs (see Methods). This is the first study to measure microglia calcium activity over all three phases after laser burn and throughout the neuroinflammatory response to infection.

**Fig. 4.**
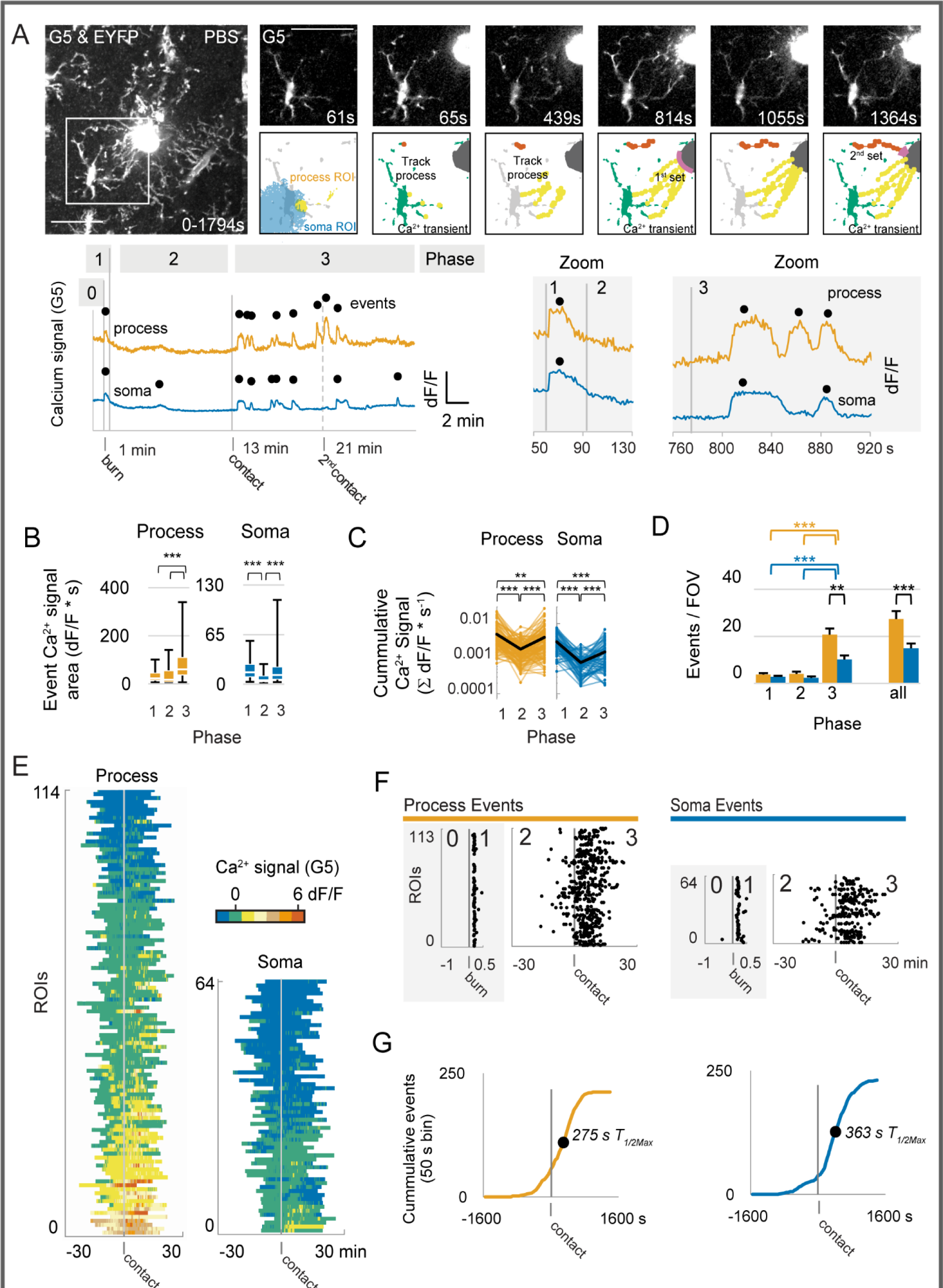
Large amplitude and frequent calcium events occur shortly after a laser burn and when processes contact burned region in slices from PBS treated mice. Microglia processes detect laser burn damage in an acute brain slice and then move toward the central burn region in PBS-injected mice. **(A)** The response phases included 1 minute of baseline and a high-power laser burn to the central region of an acute brain slice. Microglia respond to this damage with an immediate intracellular calcium event in the G5 dF/F trace from both process and soma regions, microglia processes then grow toward the burn, and come into close contact with the burn during the 30 min imaging period. **(B)** Median individual event signal areas (t-test) and **(C)** cumulative time-normalized calcium signal for individual ROIs over the response phases (Wilcoxon signed-rank test). **(D)** The number of events per FOV grouped for each phase and all phases together. **(E)** Heatmap of calcium signal and **(F,G)** event distribution displays that most calcium activity occurs immediately after the burn and again when processes come into close contact with the burn lesion (t-test). *p<0.05, **p<0.01, ***p<0.001. Mice n= 8, images n=17. Scale bar 50 µm.

**Fig. 5.**
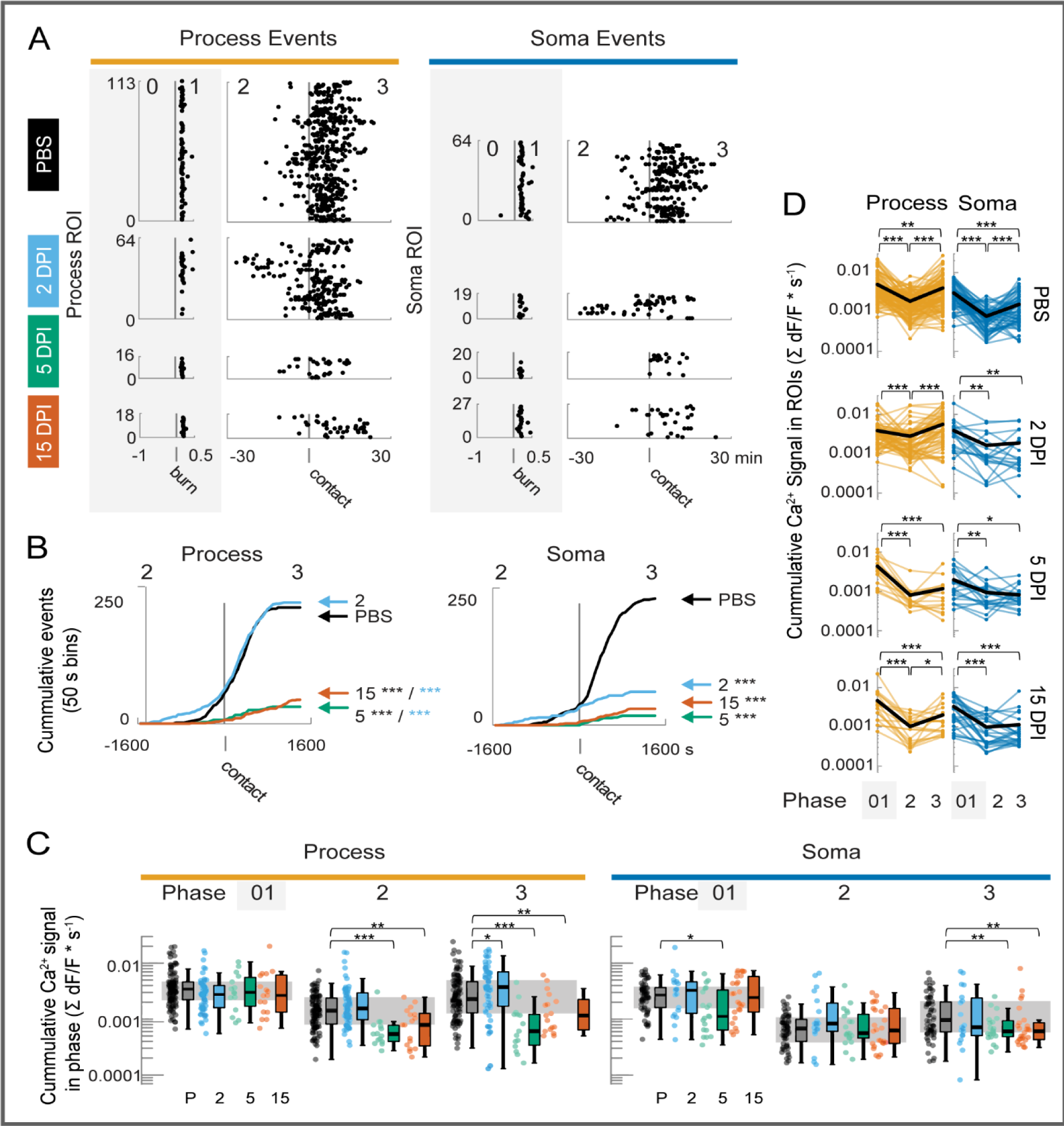
After viral brain infection, microglia respond to new damage signals with fewer intracellular calcium events at 5 and 15 DPI. **(A)** Calcium events (black dots) in microglia ROIs increase in frequency closer to the time when growing processes contact the burn area. Microglia subcellular locations (soma vs. process) display different response frequency when interacting with burn damage. **(B)** At 2 DPI TMEV, the cumulative frequency distribution in process ROIs is similar to PBS, while soma ROIs have lower event frequency (Kolmogorov-Smirnov test). And **(C)** comparison of cumulative calcium signal across viral infection groups (Mann-Whitney test). **(D)** Cumulative time-normalized calcium signal for individual ROIs over the response phases (Wilcoxon signed-rank test). Mice n=8, 5, 5, 6 and images n= 17, 7, 6, & 9 for PBS and TMEV 2, 5, and 15 DPI respectively. ***p<0.001, **p<0.01, *p<0.05.

Calcium transients in microglia can be readily observed in the branched cellular processes as well as in the soma. Immediately after a laser burn, <u>Phase 1</u> is characterized by a prolonged calcium event occurring throughout the process and soma ROIs (black dots in **Figure 4A**) with a duration range of 20-70 seconds. As the processes extend toward the burn in <u>Phase 2</u>, smaller amplitude calcium events occur in somas (**Figure 4B**, p<0.001). Although there was no difference in the amplitude (dF/F) of calcium events in processes moving from Phase 1 to Phase 2 (**Figure 4B**) both somas and processes displayed a reduced cumulative calcium signal at Phase 2 when normalized to time (**Figure 4C**). Both the frequency and basal level of calcium activity could influence signal transduction within microglia (Hoffmann *et al*., 2003). We therefore assessed this cumulative magnitude as the cumulative sum of the calcium signal change normalized to the time in each phase. When processes come into close contact with the burn area in <u>Phase 3</u>, calcium events have larger event signal areas per second (**Figure 4B**), a larger cumulative calcium signal in both soma and process ROIs compared to Phase 2 (**Figure 4C**) and occur at a higher frequency (**Figure 4D**). In **Figure 4A**, one microglia cell from a PBS-injected mouse extends a first set of processes (yellow tracks), then later a second process comes into close contact with the burn at 1283 s (orange tracks), and a calcium event is detected in both the soma ROI and the neighboring yellow process ROI at 1367 s. We observed repeated high-amplitude calcium events after microglia processes come into close contact with burn damage in Phase 3 (**Figure 4D-F**). Additionally, as shown in the cumulative probability histogram (**Figure 4G)**, microglia processes display increased event frequency, on average, 88 s earlier than soma regions upon contact with the burn in Phase 3 (275 s vs. 363 s, respectively).

Because calcium signaling is known to be involved in microglial process extension, which is disrupted during the peak of seizures at 5 DPI during TMEV infection (**Figure 3**), we asked whether the calcium signaling response to a laser burn is also altered following TMEV infection (**Figure 5**). Calcium events for both somas and processes were identified and analyzed following a laser burn in brain slices obtained from both PBS (same data as in Figure 4) and TMEV-infected mice (**Figure 5A and Supplementary Movie 1 _ PBS control mouse and Supplementary Movie 2 _ TMEV 6 DPI**) At 2 DPI, microglia can still respond to tissue damage in a similar manner as microglia from PBS-injected mice with calcium events occurring at a similar frequency in processes (**Figure 5B**). However, during this early infection response, microglia display less frequent calcium events in soma regions (**Figure 5B**), a greater cumulative magnitude in process ROIs for Phase 3 (**Figure 5C**), and notably no difference in the cumulative calcium signal for soma ROIs between Phase 2 and Phase 3 (**Figure 5D**). Therefore, at 2 DPI, the microglia processes remained responsive or were slightly more active after detecting laser damage, whereas microglia somas started to show signs of reduced calcium signaling.

At 5 DPI, reactive microglia display a reduced ability to detect damage debris and respond via intracellular calcium signals, including a reduction in the number of active calcium ROIs per field of view (FOV) (**Figure2C**), reduced incidence of calcium events in processes, in somas, or in both (**Figure 2B**), and a reduced cumulative frequency of events from Phase 2 to Phase 3 in both processes and somas (**Figure 5D**). Notably, at 5 or 15 DPI, the time-normalized cumulative magnitude of the calcium signal in process ROIs remained comparable to the PBS group in Phase 0 to 1 for those processes that still responded (**Figure 5C**). While fewer processes extend toward the burn damage (**Figure 3B**) at 5 and 15 days post-TMEV infection, those processes that do extend had a lower magnitude of calcium signal in Phase 2 and Phase 3 at 5 DPI (**Figure 5C**) when compared to processes in slices prepared from PBS treated mice. When looking at the calcium response within an individual ROI over time across all three Phases (**Figure 5D**), the cumulative calcium response was significantly reduced in slices from TMEV-infected mice after Phase 0-1 within either process ROIs or within soma ROIs. Overall, we observed fewer processes growing toward the laser burn and fewer calcium ROIs within the processes of microglia in slices from the 5 and 15 DPI groups.

### Activated microglia have disrupted ATP-coupled calcium signaling and reduced expression of p2ry12 following TMEV infection

Activation of purinergic receptors on microglia are coupled through g-proteins to intracellular calcium release and are important for process motility towards damage, where ATP spills from necrotic cells (Koizumi *et al*., 2007; Dissing-Olesen *et al*., 2014; Moore *et al*., 2015; Sipe *et al*., 2016; Jiang *et al*., 2017). To determine if the TMEV-induced changes in microglial response to burn damage are associated with alterations in purinergic signaling, we directly applied ATP (100 µM) to acute brain slices using a puffer pipette and measured microglia calcium responses at 5 and 15 DPI with TMEV (**Figure 6A**). Extracellular ATP is rapidly converted to ADP, AMP, and adenosine, and exogenous application of ATP can therefore assess changes in multiple types of purinergic receptors. Microglia expressing the calcium sensor GCaMP5 in slices prepared from PBS-injected mice responded to ATP application with a robust calcium transient (**Figure 6B**). Following TMEV infection, the microglial intracellular calcium transient was significantly reduced in slices obtained from the 5 DPI group compared to the PBS control group (**Figure 6C**, -0.2±0.4 dF/F*s *versus* 8.0±0.8 dF/F*s, respectively, p<0.001). In contrast, there was no significant effect of ATP application on microglia at 2 or 15 DPI compared to microglia from slices of PBS-treated mice. While the application of artificial cerebral spinal fluid (aCSF) alone did produce a small deflection in the acute brain slice, there was no intracellular calcium response to aCSF or the pressure of the puff in slices from either PBS-injected or TMEV-injected mice (**Figure 6C**). Thus, microglia display an impaired intracellular calcium response to ATP during the acute immune response and seizure period at 5 DPI and begin to display a restored capacity to respond to ATP application by 15 DPI.

**Fig. 6.**
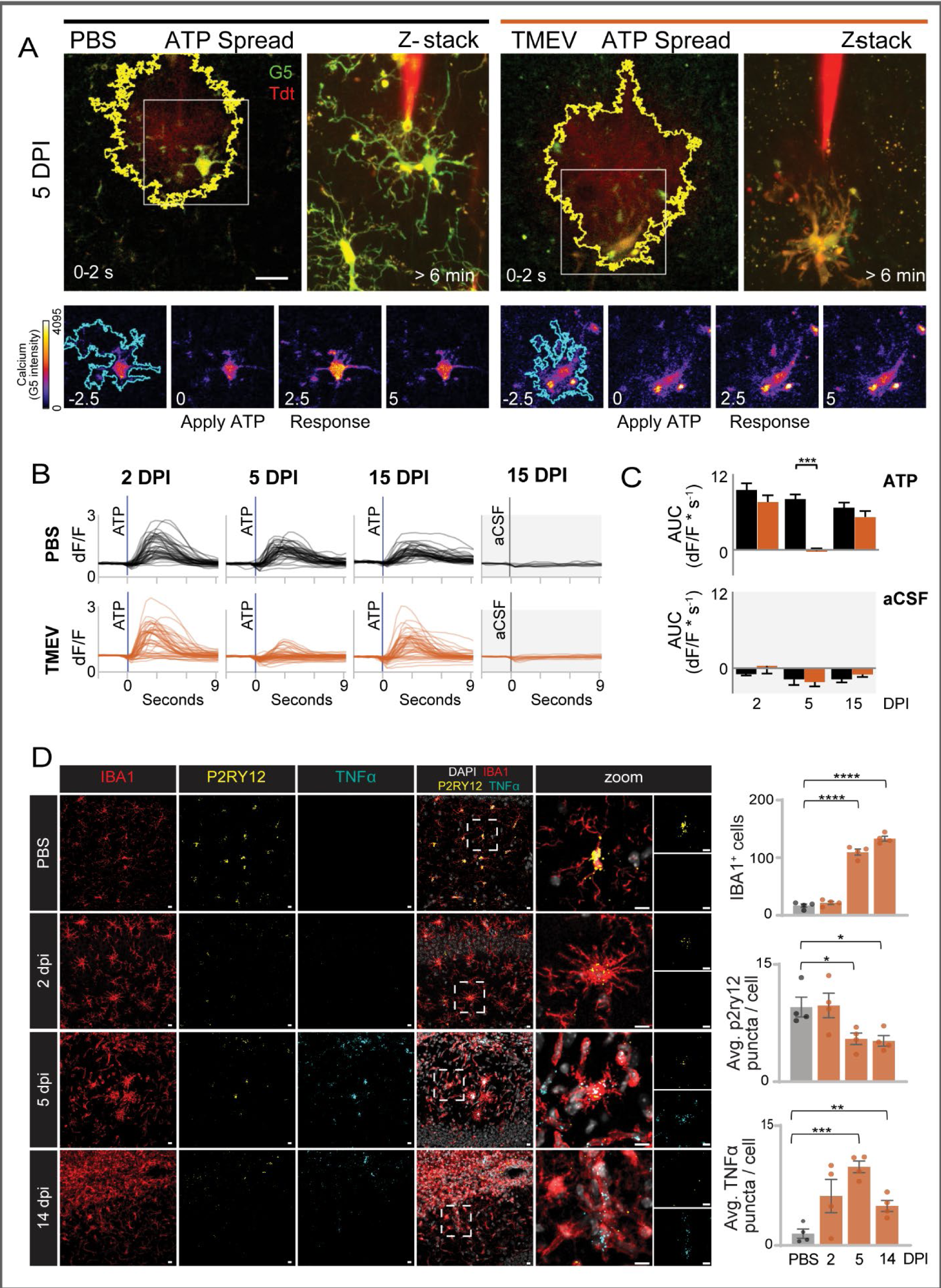
Decreased intracellular calcium response to ATP agonist and decreased *p2ry12* mRNA in microglia 5 days after TMEV infection. **(A)** GCaMP5G microglia respond an extracellular application of 100 µm ATP in acute brain slices from healthy mice. The dF/F change in intracellular calcium relative to a 4 second baseline before ATP was calculated for regions (white outlines) within the ATP spread region (Alexa568). **(B)** Calcium dF/F signal traces from microglia regions for 9 seconds after ATP application at 2, 5, and 15 DPI after PBS injection (black; mice = 9, 9, 8, cells n= 44, 46, 42) or TMEV injection (red; mice = 7, 11, 10, cells n= 47, 54, 48). The application of aCSF elicited no significant response for PBS (mice = 9, 8, 8, cells n= 9, 8, 8) and TMEV (mice n = 7, 10, 11, cells n= 7, 10, 13). **(C)** Total calcium response to ATP application was significantly reduced at 5 DPI for TMEV microglia compared to the response for PBS microglia. 2-way ANOVA with Bonferroni test (*** p<0.001). Scale bar = 20 µm. **(D)** Decreased colocalization of *p2ry12* mRNA (*in situ* hybridization) and increased *tnf-α* mRNA expression in IBA1-positive cells (IHC) in the CA1 region of the hippocampus 5 and 14 days post-TMEV infection. Mice n= 4 per group, sections = 2 per mouse (except for one 2 DPI mouse which had only one useable section). Kruskal-Wallis test for *p2ry12* (** p=0.0096). Kruskal-Wallis test for *tnf-α* (** p=0.0032). T-tests for pairwise comparisons (shown in figure) * p<0.05, ** p<0.01, *** p<0.001, **** p<0.0001. Scale bar 50 µm.

The purinergic receptor P2RY12 is robustly and selectively expressed on microglia in the CNS and is responsible for the majority of microglia process extension via Gα_s_-mediated calcium signals after ATP/ADP application or focal laser burns (Haynes *et al*., 2006; Dissing-Olesen *et al*., 2014; Eyo *et al*., 2014). *P2ry12* gene expression is moderately downregulated during acute inflammatory states, although different levels of P2RY12 expression were noted in sub-populations of activated microglia in Alzheimer’s patients (Walker *et al*., 2020). To determine *p2ry12* expression in the CA1 region of hippocampus, we performed immunohistochemistry for IBA1 and quantitated puncta of *p2ry12* and *tnf-α* mRNA (RNAscope *in situ* hybridization and Imaris) at 2, 5, and 14 DPI after TMEV-infection and compared to PBS-injected control mice. The number of IBA1-positive cells was increased at 5 and 14 DPI which could be due to both microglia proliferation (Loewen *et al*., 2016; Bell, Wallis and Wilcox, 2020) and macrophage infiltration (DePaula-Silva *et al*., 2018). The number of *p2ry12* mRNA puncta in IBA1-positive cells was significantly decreased at 5 DPI (**Figure 6D**; 5.5±1.5, p=0.04) and remained decreased at 14 DPI (**Figure 6D**; 5.2±1.3, p=0.03) compared to those in sections from PBS-injected mice (**Figure 6D**; 9.5±1.5). TNF-α is a key pro-inflammatory cytokine produced by reactive microglia (Cusick *et al*., 2013), and along with other cytokines, induces long-term synaptic scaling and promotes seizure development following TMEV brain infection (Kirkman *et al*., 2010; Patel *et al*., 2017). The number of *tnf-α* mRNA puncta within IBA1-positive cells increased significantly at 5 DPI (**Figure 6D**;9.8±1.5, p=0.0001) and at 14 DPI (4.9±1.4, p=0.0095) compared to PBS (**Figure 6D**; 1.5±1.2), which is consistent with previous hippocampal *tnf-α* mRNA and protein quantitation following TMEV brain infection (Patel *et al*., 2017; DePaula-Silva *et al*., 2019). Previously, infiltrating macrophages identified by flow cytometry increased by 5-fold at 7 DPI following TMEV-infection and returned to normal levels by 10 DPI (DePaula-Silva *et al*., 2018). This reduction in macrophages by 10 DPI would suggest the depressed *p2ry12* and elevated *tnf-α* levels are due to differences in microglia expression at 14 DPI. Taken together, these data suggest that reduced expression of *p2ry12* contributes to deficits in microglia motility and calcium signaling in response to laser burn damage and exogenous ATP application.

## Discussion

Microglia are highly adaptable innate immune cells that rapidly detect CNS pathogens and cell damage. Often, this involves acquisition of a reactive phenotype, and includes the production of defensive cytokines, including TNF-α. These reactivate pro-inflammatory microglia are known to change expression patterns for receptors and proteins involved in sensing endogenous ligands, damage signals, and pathogens. However, the present study provides strong evidence that reactive microglia following TMEV infection have altered sensitivity to damage signals.

### Main conclusions

In the present study, microglia expressing the genetically encoded calcium sensor GCaMP5G were imaged in acute hippocampal brain slices using 2-photon microscopy. This is the first functional imaging study to be performed on activated microglia in the seizure foci of TMEV-infected mice, and this adds to the growing body of microglia functional changes observed in disease-specific models. Microglia have previously been reported to display a hypertrophic morphology following TMEV infection (Kirkman *et al*., 2010; Loewen *et al*., 2016; Bell, Wallis and Wilcox, 2020), and we found a significantly reduced 3D branching network for microglia at 5 and 14 days after TMEV brain infection in live tissue. This suggests that microglia may have less surveillance capacity, with fewer processes interacting with neurons in the days and weeks following viral brain infection. Next, the functional capacity of reactive microglia to monitor their territory and to respond to damage was assessed by measuring intracellular calcium events with GCaMP5G in acutely prepared brain slices using 2-photon imaging. Microglia monitoring the hippocampal environment displayed a low frequency of spontaneous calcium events in slices obtained from PBS-treated mice and a decreased frequency at 5 and 15 DPI in TMEV treated mice. Similarly, 2 days after brain infection, microglia processes retained the ability to detect the damage produced by laser burn, measured by their ability to respond in frequency and magnitude of calcium events and elongate processes in the direction of damage. At this early phase of the innate immune response, the somas of microglia had less frequent calcium events, but the overall cumulative magnitude of the calcium signal was still comparable to microglia from PBS-treated mice. This is one of the first observations that microglia subcellular regions can display different calcium transient frequencies during inflammatory events and possibly these subcellular locations could be involved in different actions, such as actin cytoskeleton reorganization and transcriptional regulation (Färber and Kettenmann, 2006; Umpierre *et al*., 2020). However, by 5 and 15 days after brain infection, microglia had a dramatic decrease in the frequency and magnitude of both somatic and process calcium signals, and less extension of microglia processes toward the laser burn area in *ex vivo* acute brain slices. The P2YR12 receptor is thought to be responsible for a large portion of native damage responses. To test for intact purinergic signaling pathways, ATP was applied to the acute brain slices, and we found deficits in microglial ATP-induced calcium transients by 5 DPI, and which were restored by 15 days following TMEV infection. However, *p2ry12* gene expression in Iba1-positive cells in the hippocampus was significantly reduced at both 5 and 15 DPI. Therefore, the restored response to exogenously applied ATP at 15 DPI may be due to compensation from additional purinergic receptors. However, we still found motility and calcium deficits in the response to laser burn at 15 DPI, suggesting that the reduced expression of *p2ry12* may underlie process migration deficits at this later timepoint.

### Purinergic receptors and other damage detection pathways

After TMEV brain infection, there is extensive necrosis in the hippocampal CA1 neurons coinciding with the seizure peak at 5-6 DPI (Loewen *et al*., 2016; Patel *et al*., 2017). The P2y12 receptor has been regarded as a major microglia damage detection pathway, but reactive microglia show greatly reduced P2y12 expression (Haynes *et al*., 2006; Swiatkowski *et al*., 2016). In addition, several other purinergic receptors also contribute to microglial process surveillance and movement, including P2y1, P2y6, P2y13, P2×7, and A_1_ and A_2A_ receptors (Haynes *et al*., 2006; Orellana, Montero and von Bernhardi, 2013; Eyo *et al*., 2014; Calovi, Mut-Arbona and Sperlágh, 2019; Milior *et al*., 2020). In the TMEV-model, DePaula-Silva *et al*. reported significant gene expression changes for reactive microglia at four to six days after TMEV brain infection for many purinergic, G-protein coupled receptors which could compensate for decreased *P2ry12* (Gα_s_) including increased *P2ry2* (Gα_i_), increased *P2ry6* (Gα_11/q_), and increased *A2A* (Gα_s_) (DePaula-Silva *et al*., 2019). However, our current study suggests these upregulated receptors did not sufficiently compensate to maintain damage sensitivity through calcium pathways in reactive microglia at 5 days after infection. While ATP-evoked calcium responses were restored by 15 DPI, *p2ry12* gene expression, measured with *in situ* mRNA hybridization was still depressed at 15 DPI. However, the laser burn-evoked calcium responses and process extension was diminished at both 5 and 15 DPI, suggesting that non-purinergic signaling pathways may also be contributing to the long-term reduction in microglia damage sensitivity. Non-purinergic pathways involved in the microglia response to laser damage have yet to be identified, and could potentially include glutamate receptors, adrenergic receptors, cell debris/DAMPS receptors, additional extracellular matrix receptors, and complement receptors (Färber and Kettenmann, 2006).

### Subcellular calcium domains with differences in activity after brain infection

This is one of the first reports of microglia sub-cellular calcium domains in a disease condition. Over the days and weeks following viral brain infection, we identified reduced calcium activity first in microglia soma regions at 2 days after infection, followed by reduced activity in both soma and process regions at 5 and 15 days after infection. In cultured microglia, calcium domains were identified along leading edges of processes, and the calcium baseline concentration and frequency of calcium events influenced gene expression of pro-inflammatory pathways (Hoffmann *et al*., 2003; Heo *et al*., 2015; Korvers *et al*., 2016). Chronic *in vivo* microglia imaging in the kainic acid seizure model found increased somatic and process calcium activity shortly after the first seizure using a grid ROI approach (Umpierre *et al*., 2020). Long-term assessments of microglia phenotype have evaluated either whole cell or somatic regions and have not discriminated between process and somatic calcium activity. Systemic LPS injection *in vivo* leads to increased somatic calcium activity in early hours prior to morphological hypertrophy, and later, microglia had depressed somatic calcium activity at 24-30 hrs (Pozner *et al*., 2015; Riester *et al*., 2020). To accommodate growing and moving microglia processes, we used the Suite2p pipeline tailored to GCAMP5G kinetics and were able to visualize multiple calcium regions along individual or sets of processes, thus revealing differences between these two cellular compartments following infection.

### Microglia phenotype heterogeneity

Microglia that are physically closer to acute or chronic damage also display a variety of reactive phenotypes (Holtman *et al*., 2015; Dzyubenko *et al*., 2018; Bonham, Sirkis and Yokoyama, 2019; Kluge *et al*., 2019; Walker *et al*., 2020; Stoyanov *et al*., 2021). We investigated microglia function in the hippocampus near pyramidal neurons infected by TMEV, as microglia in this location are exposed to viral particles, cell debris, cytokines, and ROS (Bhuyan *et al*., 2015; Patel *et al*., 2017; DePaula-Silva *et al*., 2018). During this study, we also detected a possible heterogenous microglia phenotype in the burn response at 5 and 15 DPI. Some microglia at 5 and 15 DPI were completely non-responsive, while others could still respond with a robust calcium transient to the initial *Phase 1* burn damage but had impaired or reduced calcium response frequency after that initial response in *Phase 2* and *Phase 3*. Indeed, microglia in other brain regions may be exposed to fewer inflammatory triggers. We know microglia in the cortex retain a reactive morphology throughout 14 DPI (Loewen *et al*., 2016; Bell, Wallis and Wilcox, 2020) and it would be valuable to understand whether microglia throughout the brain remain functionally reactive and contribute to epileptogenesis in this model.

### Potential therapeutic targets at key immune transition steps during seizure development

Microglia are essential in initiating the immune response in the brain following viral infections of the CNS. Their role is complemented by infiltration of peripheral macrophages, while the later arrival of infiltrating lymphocytes eventually decreases the immune response (DePaula-Silva *et al*., 2018). In the TMEV model of TLE, mice go on to gradually develop chronic seizures in the weeks and months after the initial infection. Clearly defining when, how, and why microglia are actively responding to damage signals could help identify what immunomodulatory strategies would be most successful in preventing the development of epilepsy following infection. Our work suggests microglia are still actively sensing and responding to damage at 2 days after brain infection, and this time frame would be appropriate for therapeutic strategies to reduce microglia activation and possibly decrease the incidence of acute seizures. By 5 days after infection, microglia have a reduced capacity to respond to new damage signals due to reduced engagement of intracellular calcium transients that are coupled to both process movement and production of cytokines (Patel *et al*., 2017; DePaula-Silva *et al*., 2019). At this phase, therapeutic strategies to prevent seizures could attempt to block cytokines, and to accelerate a return of microglia to normal homeostatic roles. After this acute phase of TMEV-induced infection, seizures resolve, but following a latent period, spontaneous seizures later develop. In other chronic disease conditions, microglia never return to their normal homeostatic roles, but remain primed or chronically reactive during neurodegeneration (Brawek and Garaschuk, 2014; Holtman *et al*., 2015). Whether microglia contribute to the strengthening of subsequent seizure networks as TLE develops over time remains to be determined. Nevertheless, the present work demonstrates that a better understanding of the fundamental interactions between microglia and their environment will allow us to identify new ways to intervene in neuroinflammatory conditions.

## Materials and methods

### Animals

#### Mice

B6;129S6-Polr2a^Tn(pb-CAG-GCaMP5g,-tdTomato)Tvrd^/J(GCaMP5G) and B6.129P2(Cg)-Cx3cr1 ^tm2.1(cre/ERT2)Litt^/WganJ, (Cx3cr1-EYFP-CerERT2) were purchased from Jackson Laboratory (JAX 024477 and 021160). Male and female mice were bred to heterozygous expression of Cx3CR1-CreERT2 and GCaMP5G on a C57BL/6J background. All experiments conformed to the standards of the National Institutes of Health Guide for the Care and Use of Laboratory Animals and were approved by the University of Utah’s Institutional Animal Care and Use Committee (IACUC). Mice were provided food and water ad libitum and were maintained on a 12 h light/dark cycle in temperature- and humidity-controlled rooms.

#### Tamoxifen induced recombination

Mice 4-6 weeks old were given three doses 150 mg/kg or 200 mg/kg (i.p) tamoxifen (TAM) (Sigma-Aldrich T5648) dissolved in peanut oil (20 mg/ml) (Spectrum hi-oleic unrefined or refined) every other day to allow Cre-mediated expression of the calcium senor GCaMP5G and red cytosolic TdTomato (TdT) in Cx3cr1 expressing microglia and macrophages. The Cx3cr1 mouse strain includes a non-inducible yellow EYFP fluorophore. After the third TAM administration, a minimum of 35 days elapsed before experiments to allow for a newly born population of bone derived macrophages to differentiate and not express the TAM-induced fluorophores (Parkhurst *et al*., 2013). Thus, in these experiments, infiltrating macrophages are identified by yellow EYFP expression, while microglia express yellow EYFP, calcium-sensitive green GCaMP5G, and red TdT.

#### TMEV infection and seizure observations

Male and female mice were anesthetized using a mixture of isoflurane and compressed air. Mice were infected by delivering 2×10^5^ plaque forming units (pfu) of the Daniels strain of TMEV intracortical injection to a depth of 2 mm in the temporal region of the right hemisphere (posterior and medial of the right eye) (Libbey *et al*., 2008). Control mice were injected with 20 µL sterile PBS instead of virus. Mice were video recorded while their cage was briefly agitated, and they were monitored for seizures twice daily between days 3 and 7 DPI. The intensity of the seizure activity was graded from the video recording on a modified Racine scale: stage 3, forelimb clonus; stage 4, rearing; stage 5, rearing and falling; and stage 6, clonic running or jumping around the cage. Mice at 2 DPI did not have seizures during handling but did have mild weight loss. Mice at 5 and 15 DPI were used for brain slice preparation if they displayed at least one grade 3 seizure during the monitoring period.

#### Acute brain slice preparation

Acute brain slices were prepared from animals at 8-14 weeks of age and at 2 DPI, 5 DPI (4-6 days after either PBS or TMEV injection), and 15 DPI (14-16 days after injection). Mice were deeply anesthetized with isoflurane and were non-responsive to a foot pinch prior to decapitation. Coronal sections containing the hippocampus (400 µm) were cut on a vibratome (Vibratome 3000, Vibratome Company) using an ice-cold sucrose solution: 200 mM sucrose, 3 mM KCl, 1.4 mM NaH_2_PO_4_, 26 mM NaHCO3, 10 mM Glucose, 2 mM MgSO_4_, and 2 mM CaCl_2_ (osmolarity: 290-300 mOsm). Sections were transferred to a recovery chamber containing room-temperature aCSF: 126 mM NaCl, 2.5 mM KCl, 1 mM NaH_2_PO_4_, 26 mM NaHCO_3_, 10.5 mM Glucose, 1.3 mM MgSO_4_, and 2 mM CaCl_2_ (osmolarity: 307-311 mOsm). Sections were given a minimum of 1 h to recover before two-photon imaging and were imaged for a maximum of up to 5 hrs after slicing. All solutions were bubbled with 95% O_2_/5% CO_2_ and titrated to a pH of 7.35-7.4. Reagents used to make solutions were purchased from Sigma-Aldrich.

#### Structural and calcium imaging

Two-photon (2-P) calcium imaging and structural imaging was performed on a Prairie Ultima system (Bruker Corporation) using a Mai Tai DeepSee EHP 1040 laser (Spectra Physics) at 69 mW laser power, Prairie View software, a 20X water-immersion lens (NA: 0.95, Olympus), and emission bandpass filter at 560 nm to split green from red wavelengths (Bruker 370A510816). GCaMP5G and EYFP contribute to the majority of the green channel signal, while TdT contributes to the majority of the red channel signal. Brain sections were continuously perfused with aCSF at a rate of 1-3 mL/min by a peristaltic pump system. Bath temperature was maintained at 28-32 °C by an in-line heater (TC-324C, Warner Instruments). Microglia expressing GCaMP5G and TdT were imaged in both the left and right hippocampal CA1 regions of *stratum oriens, stratum pyramidale,* and *stratum radiatum* at a depth of approximately 60-80 µm from the surface of the slice to avoid superficial areas damaged by the slicing procedure.

Images of microglia morphology were acquired at 1000 nm excitation where red TdT is more optimally excited and green GCaMP5 is still visible. Z-stacks of 81 x 1 µm images were acquired beginning at 40 µm from the surface using 9.2 µs/pixel dwell, 1024 x 1024 pixels per frame, 2x optical zoom (292 x 292 µm field of view (FOV)), and 460 pockels laser power.

Calcium imaging of spontaneous transients was performed at 920 nm excitation wavelength for GCaMP5G. Time series were acquired at 0.93 Hz, 3.2 µs/pixel dwell, 512 x 512 pixels per frame, 2x optical zoom (292 x 292 µm FOV), and 290 pockels laser power for 15 min.

For ATP-agonist calcium imaging, stock 10 mM ATP (Tocris 3245) in reverse osmosis water was stored for up to 1 month at -80 °C. Daily working solutions were diluted to 100 µM ATP in aCSF with 15 µg/mL Alexa568 (Invitrogen A33081) to visualize the puff region. Puff pipettes were pulled by a HEKA PIP 6 electrode puller from 1.5 mm OD, thin-walled borosilicate glass and had an open tip resistance of 2.1-3.6 MΩ. ATP was dispensed using a Picospritzer III system (Parker Instrumentation) with 6 PSI pressure for 350 ms. The ipsilateral side receiving the TMEV or PBS injection was imaged in the CA1 regions of *stratum pyramidale* and *stratum radiatum* and either ATP or aCSF was applied to different fields of view in the same brain slice. Time series images were acquired at 920 nm excitation, 2 Hz, 1.2 µs/pixel dwell, 512 x 512 pixels per frame, 4x optical zoom (146 x 146 µm FOV), and 240 pockels laser power for a 30 s baseline and 5.4 min after the puff. After the time series image was completed, a z-stack of the same region was acquired at ±15 µm with 1 µm spacing using 9.2 µs/pixel dwell, 1024 x 1024 pixels per frame, 4x optical zoom (146 x 146 µm FOV), and 460 pockels laser power.

Calcium imaging and process movement in microglia in response to a high-power laser burn was acquired for (1) a baseline period (920 nm for 63 s at 0.97 Hz, 3.2 µs/pixel dwell, 512 x 512 pixels per frame, 3x optical zoom with 195 x 195 µm FOV, and 290 pockels laser power), (2) a high-power burn on the central 12 x 12 µm region (800 nm for 2.7-5.3 s at 1.1 Hz, 3.2 µs/pixel dwell, 512 x 512 pixels per frame, 50x optical zoom with 12 x 12 µm FOV, and 660-700 pockels laser power for 2.7-5.3 s), and (3) followed by the calcium and process movement responses (920 nm for 29 min at 0.97 Hz with the same settings as baseline period). Brain sections from TMEV mice required longer burn durations and power to achieve a similar diameter burn. After the burn, a lag period of 25 s was required for the laser to return to 920 nm prior to acquiring the calcium response time series. Slices remained stationary for these extended imaging durations if the slice was firmly adhered to the supporting meshwork by vacuuming aCSF from below the slice three times, and if the slice received 30 min of room temperature post-slice incubation and 30 min at 28-32 °C prior to imaging. For each burn time series, the TdT maximum-pixel-intensity over time image was manually thresholded to mask the burn area, and the average burn diameter was 21 ± 5 µm.

### Image analysis

#### Morphology of microglia

Z-stack signal loss due to emission light scatter in deep z-slices was normalized to the most superficial z-slice using Stack Contrast Adjustment in ImageJ (Michalek, Capek and Janacek, no date). A best-fit 3D skeleton was computed inside the microglia volume using 3DMorph (York *et al*., 2018) with user-defined threshold settings on the green channel to accommodate different background fluorescent levels. The parameters reported for each cell in the FOV included the cell volume, cell territory (including cell and non-cell space), ramification index (territory/cell volume), number of branch endpoints, number of branching points, and the maximum, average and minimum branch length. One acute slice per mouse was imaged for n=7, 6, and 8 mice for the PBS control group at 2, 5, and 14 DPI, respectively, and n=6, 6, and 7 for TMEV-injected mice at 2, 5, and 14 DPI, respectively.

#### Frequency of spontaneous calcium transients following burn damage

Regions containing spontaneous calcium fluctuations were identified in a conservative manner using the fully automated Suite2p toolbox (Pachitariu et al., 2017; Stringer and Pachitariu, 2019). Suite2p identifies correlated pixels that fluctuate on a time scale appropriate for the fluorescent sensor and the imaging rate compared to a stable local background region which compensates, in an unbiased fashion, for the different background fluorescence levels in TMEV-infected and PBS control brain slices. Then a weighted pixel intensity for each active region is reported. Slice drift during the imaging period was reduced by registering relative to the TdT signal over time. Calcium signal changes were calculated relative to the average baseline for the first 50 image frames. For the dataset evaluating baseline frequency of spontaneous calcium transients, ROIs were identified using the *findpeaks* function in MATLAB (>2.5x mean prominence, >5x STD running 50 frame average dF/F, and >0.4 dF/F amplitude), and active ROIs containing active signals were confirmed by a reviewer. The number of events per FOV, events per ROI, and ROIs per FOV were normalized to the total image time.

#### Process movement tracking, classifying phases of laser burn response, and calcium event detection

The movement of microglia processes toward the burn damage zone was tracked every 60 s using ImageJ MTrackJ (Meijering, Dzyubachyk and Smal, 2012), and the process locations and velocities were recorded. The area of the burn zone was measured on the TdT maximum intensity over time image using ImageJ. The process location and velocity were used to calculate the time at which the growing process came into contact with the burn zone, which is referred to as the “*contact time*”. A reviewer classified the subcellular location of each ROI (soma *versus* process), and process events were further classified if they overlayed a process growth trajectory. For each cell, all tracks associated with unique ROIs were included, and spatially redundant track/ROI pairs removed. Calcium events were then classified in four phases of microglia response: *Phase 0* baseline from 0 to 58.2 s (0 to 60 frames); *Phase 1* initial wave of burn damage until 92.7 s (61 to 90 frames); *Phase 2* while microglia processes hone in on the burn area until the *contact time* (frame 91 until *contact time*); and *Phase 3* after calculated *contact time* (*contact time* until frame 1740). The cumulative calcium signal for each phase was calculated as Σ(F_i_ - F_i-1_)/seconds on a 0.012 Hz lowpass filtered dF/F for each ROI.

For the calcium transients that occurred in response to the laser burns, events were identified by reducing noise with a lowpass filter (0.06 Hz), the local minimums and maximums identified using the *findpeaks* function in MATLAB (local max >1.8 STD of mean prominence, local minimum is negative of the signal and <1.5 STD mean prominence), and correct event identification confirmed by a reviewer. Event amplitude, duration, and area under the curve were calculated on the unfiltered dF/F signal relative to the local minimums. The signal-to-noise ratio (SNR) was calculated as the difference of the event signal and the background signal divided by the summation of the squared variances for each. The SNR was 7.4±1.8 (mean±SEM) for all GCaMP5G events in microglia identified after laser burn in acute brain slice.

#### ATP agonist spread and calcium response

The area of ATP agonist spread in the brain slice was identified by the spread of Alexa568 in the post-application period. Regions of interest slightly larger than cells within the ATP spread area were identified using the semi-automatic GECIquant (Srinivasan et al., 2015) with user-defined thresholding settings. Image noise was reduced with a hybrid 3D median filter in ImageJ (C.P. Mauer & V. Bindokas). Calcium signal change (F-F_0_)/F_0_ was calculated from the mean pixel intensity in the ROI (F) for each point in time compared to the mean pixel intensity for baseline 4.5 s before the application (F_0_).

#### Dual RNAscope in situ hybridization and immunohistochemistry

TMEV-infected and PBS-control mice were sacrificed at 2, 5, or 14 dpi. Animals were anesthetized through intraperitoneal injection with pentobarbital, and transcardial perfusions were performed with PBS followed by 10% neutral buffered formalin solution (NBF). Brains were then postfixed for 24 hours in 10% NBF and transferred to a 15%/30% sucrose gradient for cryoprotection. Tissue was sectioned coronally to 15 µm on a freezing stage microtome (Leica, Buffalo Grove, IL). Slides were mounted with duplicate sections from each brain, and slides were immediately processed for RNAscope. Slides were also prepared for positive and negative control probes for each brain.

Fluorescent *in situ* hybridization (FISH) was performed as per the manufacturer’s instructions using RNAscope® Multiplex Fluorescent Reagent Kit v2 for Fixed Frozen Tissue using catalog probes TNF-α (Cat No. 311081) and P2RY12 (Cat No. 317601-C2). Briefly, brain tissue sections were dehydrated by 50%, 70%, and 100% ethanol gradually for five minutes, then boiled for 5min in 1X Target Retrieval Reagent. A hydrophobic barrier was applied (ImmEdge™, Vector Laboratories H-4000), and sections were incubated with Protease III Reagent for 30 minutes in a 40°C hybridization oven (Boekel Scientific, Model 136400). Probe hybridization (2hr) followed by signal amplification and development using Opal™ Dyes 690 and 520 (1:1500) steps were performed using the 40°C hybridization oven. Slides were washed 2X in Wash Buffer reagent and followed with subsequent immunohistochemical staining.

Sections were blocked in CytoQ ImmunoDiluent & Block Solution (Innovex NB307-C) containing 0.3% Tween-20 for 1 hour. Tissue was incubated overnight at 4°C with primary antibody directed to ionized calcium-binding adaptor molecule 1 (IBA1) (Novus Biologicals NB100-1028, 1:500) diluted in CytoQ containing 0.05% Tween-20. Sections were then washed five times with CytoQ containing 0.1% Tween-20 and incubated for 2 hours at room temperature with secondary antibody AlexaFluor donkey anti-goat 546. Slides were counterstained with DAPI, (Advanced Cell Diagnostics) for 30 seconds, then rinsed five times with PBS and mounted with Prolong Gold antifade reagent (Molecular Probes) using No. 1.5 coverslips.

Images were captured with a Nikon A1R confocal microscope (Nikon Instruments, Melville, NY) at the University of Utah Cell Imaging Core Facility using a x40/1.3 oil objective. Laser output, photomultiplier, and offset settings were adjusted to minimize saturated pixels and maximize contrast across samples. Once optimized, the laser settings were held constant between images acquired across all slides. Regions of interest, specifically the right dorsal CA1 region of the hippocampus were initially identified using epifluorescence in the DAPI channel. Once a region was selected, laser-scanning mode was used to acquire 6 x 1 µm z-stack optical images for each brain from duplicate sections using Nikon’s Confocal NIS-Elements Acquisition Software. Probe specificity was confirmed using negative (dapB) and positive (PPIB) control probe sections.

3D confocal stacks were first pre-processed using Fiji/ImageJ software (National Institutes of Health, Bethesda, MD). Images were processed to maximize the signal-to-noise ratio in batch using the “Hybrid 3D median Filter plug-in (Mauer and Bindokas, no date) followed by a rolling ball background subtraction. The resulting images were analyzed with Imaris software (version 5.5.0; Bitplane AG). Imaris was used to generate spots for the FISH probes in each channel with the program’s “Spots” feature. A user-defined intensity threshold was determined for each image in order to eliminate varying background intensities and minimize spurious spot identification. The experimenter thresholded such that most puncta identified outside the nuclei and IBA1^+^ cell volumes were eliminated.

IBA1^+^ cells were determined using the Imaris “Surfaces” feature. RNA “spots” that were localized within the “surfaces” were quantified and exported using the Imaris “statistics” feature.

#### Statistical analysis

Statistics were performed with GraphPad Prism 5, Microsoft excel, and the Real Statistics Resource Pack Software (Release 7.6). Copyright (2013 – 2021) Charles Zaiontz. www.real-statistics.com.

#### Illustrations

Illustrations were adapted from Servier Medical Art (https://smart.servier.com/)

## Acknowledgements

We thank E. Jill Dahle for technical support and Drs. Anthony Umpierre and Ana Beatriz DePaula-Silva for helpful discussions. We thank the University of Utah Cell Imaging Core Facility for use of the Nikon A1R confocal microscope and IMARIS analysis platform.

## CRediT Author Statement

Glenna J. Wallis (conceptualization, methodology, validation, formal analysis, investigation, data curation, writing-original draft, writing-review and editing, visualization, funding acquisition); Laura A .Bell (conceptualization, methodology, investigation, validation, visualization, formal analysis, data curation, writing-original draft, writing – review and editing, funding acquisition); Lauren Buxton (methodology, investigation, data curation); John Wagner (software, data curation); Lakshmini Balachandar (writing -review and editing); Karen S. Wilcox (conceptualization, methodology, writing-review and editing, supervision, project administration, funding acquisition).

## Abbreviations

2-P: two-photon
3D: three dimensional
A1: A1 adenosine receptor
A2A: A2A adenosine receptor
aCSF: artificial cerebrospinal fluid
ADP: adenosine diphosphate
AMP: adenosine monophosphate
ATP: adenosine triphosphate
CA1 & CA3: subfields 1 and 3, respectively, of the cornu ammonis region of the hippocampus
CNS: central nervous system
CreERT2: Cre recombinase – estrogen receptor T2
Cx3cr1: C-X3-C Motif Chemokine Receptor 1
DAMP: damage-associated molecular pattern
dF/F or ΔF/F: the change in fluorescence intensity relative to the baseline fluorescence intensity
DPI: days post-infection
EYFP: enhanced yellow fluorescent protein
FISH: fluorescent *in situ* mRNA hybridization
FOV: field of view
G5: genetically encoded green calcium indicator variant 5G
GCaMP: genetically encoded calcium indicator
Hrs: hours
Hz: hertz
IBA1: ionized calcium-binding adapter molecule 1
IL-6: Interleukin 6 cytokine
i.p.: intraperitoneal injection
mOsm: milliosmole
Min: minute(s)
mL: milliliter
mm: millimeter
mM: millimolar
mRNA: messenger ribonucleic acid
ms: millisecond
mW: milliwatt
MΩ: megaohm
NA: numerical aperture
NADH: nicotinamide adenine dinucleotide + hydrogen
NBF: neutral buffered formalin
NG2: Nerve/glial antigen 2, also known as chondroitin sulfate proteoglycan 4 (CSPG4)
nm: nanometer
OD: outer diameter
P2RY12: purinergic receptor P2Y12
P2RY: P2Y purinergic receptors
PBS: phosphate-buffered saline
PC: *Polr2a* gene
Pfu: plaque-forming units
PSI: pounds per square inch
RNA: ribonucleic acid
ROI: region of interest
ROS: reactive oxygen species s second(s)
SEM: standard error of the mean
SNR: signal-to-noise ratio
STD: standard deviation
TAM: tamoxifen
TdT: tdTomato fluorescent protein
TLE: temporal lobe epilepsy
TMEV: Theiler’s murine encephalomyelitis virus
TNF-α: tumor necrosis factor alpha
µg: microgram
µL: microliter
µm: micron (also known as micrometer)
µM: micromolar
µs: microsecond.

